# Bottlenecks and Resistance Evolution against an Antimicrobial Peptide

**DOI:** 10.1101/2025.03.27.645738

**Authors:** Onur Erk Kavlak, Luisa Linke, Elisa Bittermann, Mathias Franz, Jens Rolff

## Abstract

Bottlenecks – a severe reduction in population size—are ubiquitous for bacteria in natural systems. For example, during the complete metamorphosis of insects, the gut microbiota is bottlenecked due to a massive secretion of immune effectors such as antimicrobial peptides (AMPs) into the gut. However, the effect of natural bottlenecks on resistance evolution of bacteria to AMPs is currently unknown. Here we measured the bottlenecking of the gut microbiota of *G. mellonella*, the greater wax moth. Based on these population estimates, we tested *in vitro* the influence of population size on the adaptation of *E. coli* against an AMP. We used wild-type *E. coli* and mutator strains with a 100-fold increase in mutation rates to partly disentangle population size from mutant supply. We found that large *E. coli* mutator strain populations evolved higher resistance than small populations. Population size, however, did not affect adaptation in wild-type strains. They were not able to evolve resistance under our experimental conditions. This shows that natural bottlenecks can benefit insect hosts by combating the evolution of AMP resistance to their resident gut microbiota, but this should also be applicable to infection in other organisms that use AMPs as an immune defense.

## INTRODUCTION

Bottlenecks – a drastic decline in population size – are common in natural systems (1). During complete metamorphosis in insects, for example, more than 60% of all animal species are insects with complete metamorphosis (2), bacterial populations are bottlenecked due to a combination of high concentrations of antimicrobial peptides (AMPs) and lysozyme in the metamorphic gut (3,4). Bottlenecks can influence the evolution of microorganisms by altering the emergence and fixation of new beneficial traits while mutations provide the raw material that selection acts on. (5). However, how bottlenecks and the mutation rate affect adaptation towards immune effectors such as antimicrobial peptides has received little attention.

AMPs are crucial immune effectors to control microbes (6). Recent research on *Drosophila melanogaster* has revealed that fruit flies synthesize AMPs to purge pathogenic microbes and control the symbiont abundance that otherwise can compete for resources with the host (7). In the Lepidopteran insect *Galleria* mellonella, the greater wax moth, a variety of antimicrobial peptides are synthesized and secreted into the gut before gut renewal during metamorphosis (8). The combination of these antimicrobial effectors, mainly AMPs and the antimicrobial enzyme lysozyme, significantly decreases the resident gut microbiota, pathogenic bacteria are cleared from the gut but symbionts are maintained (9).

Bottlenecks can have significant consequences for the adaptation of bacterial populations (10– 12). They change the fixation probability of a beneficial mutation as well as the ability to adapt to stressful conditions such as antimicrobial drugs (5,13). Theory suggests that larger bottleneck sizes, increase the adaptation by lowering the probability of beneficial mutations getting lost (1,5). Similarly, small bottleneck sizes may decrease the adaptation due to weaker selection and stronger genetic drift causing the accumulation of deleterious mutations (2,13). An experimental study on *E. coli* showed that larger bottleneck sizes result in acquiring different resistance mutations against a fluoroquinolone antibiotic compared to smaller bottleneck sizes (14). Studies using different microorganisms, bacteria, unicellular green alga, and different stressors such as temperature, salt, or antibiotics also found that adaptation to the stressors is more likely under large bottlenecks. (13,15–17), but see (10,12). Therefore, the strength of a bottleneck can significantly determine the resulting adaptation.

In addition to bottlenecks, mutations affect drug adaptation. For instance, mutator phenotypes with higher mutation rates can emerge in natural bacterial populations (18). Higher mutation rates contribute to acquiring rare beneficial mutations faster, allowing them to respond to selection much quicker (19). For example, *Perron et al*. 2006 reported that mutator strains of *P. aeruginosa* under selection with the AMP pexiganan evolved higher resistance than wild type strains. For *E. coli*, they found the opposite pattern (20).

Here, we investigated the effect of natural bottlenecks occurring during insect metamorphosis on the evolution of resistance. We first measured the population size changes of the gut microbiota throughout the complete metamorphosis of the wax moth, *G. mellonella*, with culture-based methods. This provided us with a range of population sizes during insect metamorphosis that informed the bacterial population sizes in our experiments. Using wild-type(wt) *E. coli* MG1655 and its corresponding mutator strain (*mutS*) under different bottleneck sizes allows to at least partially uncouple the effect of population size from mutant supply on the evolution of resistance.

## MATERIAL AND METHODS

### Insect rearing

*G. mellonella* larvae were purchased from a commercial seller for pet food (fauna topics, Mahrbach, Germany) and reared in the dark at 25°C. The insect diet was optimized from (21) which consisted of different grains (wheat bran, oat bran, and rice bran), honey, glycerol, and bee wax.

### Gut microbiota counting

We used culture-based techniques to count symbionts (*Enterococcus spp*.) in the gut because the symbionts were culturable on standard microbial media (22,23). We first dissected the *G. mellonella* guts at each of the five developmental stages, as described in (24). Briefly, **stage 0** is a wandering and feeding larva. **Stage 1** is a wandering larva that has stopped feeding and started spinning. Pigments in stemmata have not started to migrate and the midgut is empty. **Stage 3** is a later cocoon-spinning larva, in which the pigments have migrated from the stemmata but are still in contact with the cuticle. The larval gut epithelium has completely detached and drops in the lumen. **Stage 5** is the prepupa that does not spin and stemmata pigments are invisible. **Stage A** is adults which emerged from pupa. Gut samples from each developmental stage (number of gut samples = 20,26,21,19,3 for stages 0,1,3,5, A respectively) were suspended in 200 µl of PBS buffer. Individual guts were homogenized at 30 Hz for 20 seconds with a tissue homogenizer (Mixer Mill, Retsch GmbH, Germany). Then gut homogenate was serially diluted and placed onto tyriptic soy agar (TSA) at 37 °C overnight. Lastly, we counted the colonies to assess colony-forming units (CFUs).

### Bacterial strains and growth conditions

Due to their well-known biology and genetic repertoire, we used *E. coli* MG1655 bacteria as a proxy for gut microbiota. Experiments were done by using a wild-type (wt) strain and a corresponding mutator strain *mutS* (∼100 times higher mutation rate than wild-type) that has a defected methyl-mismatch system (20).

### Selection Experiment

Before starting the selection experiment, we first preadapted *mutS* and wild-type strains in Müller Hinton Broth (MHB) media for a week. Briefly, we diluted the bacterial cultures 1:1000 and incubated them in 50 ml of Falcon tubes containing 3.7 ml of MHB at 37 °C, 200 rpm. Then we used six bacterial cultures derived from a single colony for wild-type and *mutS* strains. Selection lines were exposed to pexiganan (GIGKFLKKAKKFGKAFVKILKK-NH_2_, Innovagen-Sweden) treatment, and positive control lines passaged in MHB. The evolution of resistance to pexiganan was conducted with two different bottleneck sizes: small (≈10^5^ CFU) and large (≈10^7^ CFU). In this way, we mimicked the gut microbiota abundance at the bottleneck (Fig. 1). The selection was started with 1 µg/ml of pexiganan concentration, which equals half of the minimum inhibitory concentration (½ MIC) of *E*.*coli*, and was doubled every four days. For small bottleneck ≈10^5^ and large bottleneck ≈10^7^ CFU were inoculated into fresh MHB at the end of each growth cycle (every ≈20 hours). The inoculum size did not exceed 10% of the total culture volume (200 µl) to ensure that all populations have sufficient nutrients to grow. Small and large transfer bottlenecks grew up to the same carrying capacity (≈10^9^ CFU) at the end of each growth cycle in a 96 well plate. We selected cultures for forty days or until they went extinct. The last surviving cultures under the pexiganan treatment are called endpoint cultures. Extinction was described as two days without growth in the 96-well plates (an equal OD with the blank) and no growth on a Müller Hinton plate. Fossil records from each growth cycle were preserved in 25% glycerol stocks at -70 °C.

**Figure 1.**
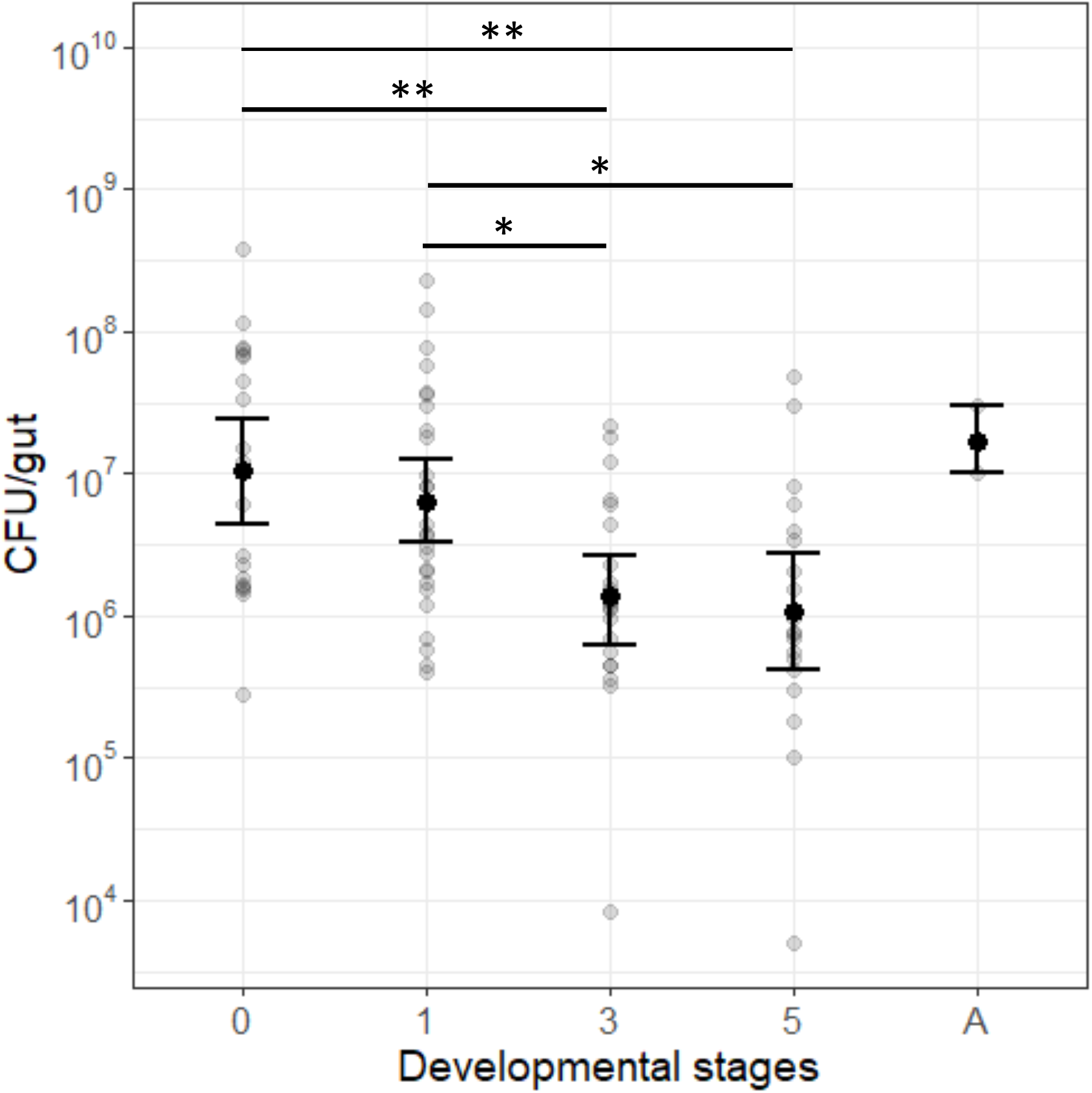
The bottleneck in gut microbiota during metamorphosis. Numbers indicate the stages and “A” adult. Each dot shows one gut sample and black solid points show the mean CFU count. Error bars represent 95% confidence intervals. Star signs show statistically significant differences between developmental stages (* p < 0.05, ** p< 0.01, Table 1).

### Antimicrobial Susceptibility Testing

We assayed the resistance of selection lines by measuring MIC based on the European Committee on Antimicrobial Susceptibility Testing (EUCAST) guidelines using the protocol described in (25). Shortly, *E. coli* cells were grown in 3 ml of MHB at 37°C for 200 rpm overnight. Then 100 μl of 10^6^ CFU/ml were inoculated with 100 µl of two-fold serially diluted pexiganan solution ranging from 512 to 0.25 µg/µl in 96 well plates. The MIC was considered the lowest concentration that inhibited visible bacterial growth after ∼20 hours of incubation. We used 3 biological replicates per AMP concentration. Since EUCAST does not provide breakpoint concentrations for the AMP-resistant isolates, we described the evolution of resistance as the significant increase of MIC in selection lines compared to their ancestors.

### Inoculum effect

As our selection experiment used two different inoculum sizes (10^5^ and 10^7^ CFU), we checked the MIC values for these sizes for wild type and mutS strains to control for the inoculum effect (Reference). We prepared different inoculum sizes of *mutS* and wt strains. First, we inoculated the bacteria culture overnight at 37 °C, 200 rpm. Then approximate cell numbers ranging from 10^2^ to 10^8^ CFU in 200 µl of MHB (5 × 10^2^ to 5 × 10^8^ CFU/ml) were calculated using a plate reader. Two-fold dilutions of pexiganan solution (from 0.25 to 512µg/ml) were prepared in a 96-well plate. Bacterial suspensions were added to the pexiganan solutions. MIC was determined for different inoculum sizes after an incubation at 37 °C for 20 hours. 3 replicates were used to measure MIC.

## RESULTS

### Bottlenecks in gut microbiota during metamorphosis of *G. mellonella*

We measured the abundance of the gut microbiota during metamorphosis to determine bottleneck sizes in *G. mellonella*. CFU counts significantly differed among metamorphic stages (*F*_*4,89*_ = 6.236, p < 0.001, ANOVA, Figure 1). As expected, gut microbiota abundance was higher at the beginning of metamorphosis (stage 0). Towards gut replacement (stage 3), it started to decline and reached the lowest point at stage 5 and increased again at the adult stage. At the bottleneck (stage 5), microbiota abundance approximately fluctuated between ≈10^5^ and 10^7^ CFU/gut. Based on these results, we used 5 × 10^5^ CFU/ml and 5 × 10^7^ CFU/ml for small and large bottleneck sizes, respectively, to conduct experimental evolution (**Figure *1***).

### Survival and evolution of resistance to pexiganan

We first recorded the time to extinction for the different treatment groups. As a result, small bottlenecks of wt and *mutS* went extinct consecutively and this was followed by large bottlenecks of wt and *mutS* strains (Figure 2a). All large bottlenecks of *mutS* strain survived at the last concentration of 512 µg/ml, which equals 256 times higher MIC than their ancestor. To determine the level of adaptation after selection with pexiganan, we measured the MIC of populations at their endpoints. We found a significant interaction of strain and population size for adaptation to pexiganan (*F*_*1,24*_ =26.24, *p* < 0.001, Figure 2b). For the wild-type strain, we did not observe any resistance evolution and accordingly observed no statistically significant effect of population size (t =-1.116, *p* =0.684). In contrast, the small and large *mutS* populations evolved resistance towards pexiganan with large populations showing markedly higher resistance than small populations (on average 48.5 compared to 6.13 MIC fold increase, t = 8.36, p < 0.001).

**Figure 2.**
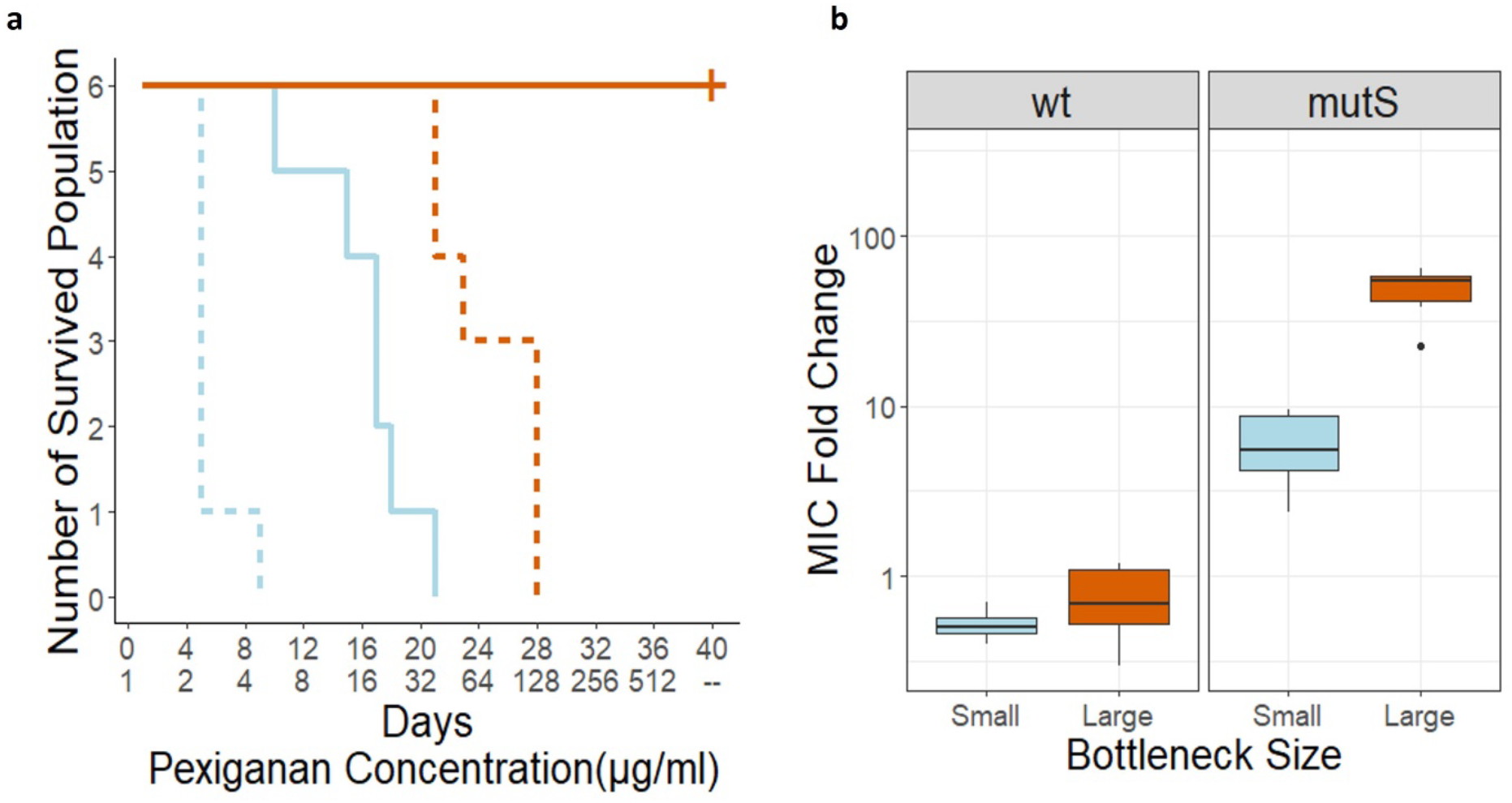
**a)** Population survival during the selection experiment. Line types show the strains and colors bottleneck size (solid lines:*mutS*, dashed lines: wt; light blue colour: small; orange colour: large bottleneck size). The first x-axis line shows the days and second x-axis corresponding pexiganan concentration at that day. The pexiganan concentration was doubled every four days. The experiment started at ½ of MIC of small populations on day 0 and was completed at 256 x MIC in day 40. **b)** Phenotypic changes of wild-type(wt) and mutator(*mutS*) strains during selection experiment. Boxplot shows the MIC fold change of treatment groups relative to their ancestor. The center line shows median and single dot indicates outlier. “ns” non-significant, ****P < 0.001; Tukey’s honest significant difference (HSD).

### The effect of inoculum size on the susceptibility to pexiganan

We tested wild-type and *mutS* strains for inoculum effect because large bottleneck populations went extinct at much higher concentrations than their actual concentrations that they adapted (Fig. 2a and Fig. 2b). We found that the increase in the inoculum size increases the MIC after a threshold (**Figure 3**). The wt and *mutS* strains had a slight increase in MIC until the population size reached 10^5^ CFU. After the 10^5^ CFU threshold, MIC increased drastically for wt and *mutS* strains (***Figure 3***).

**Figure 3.**
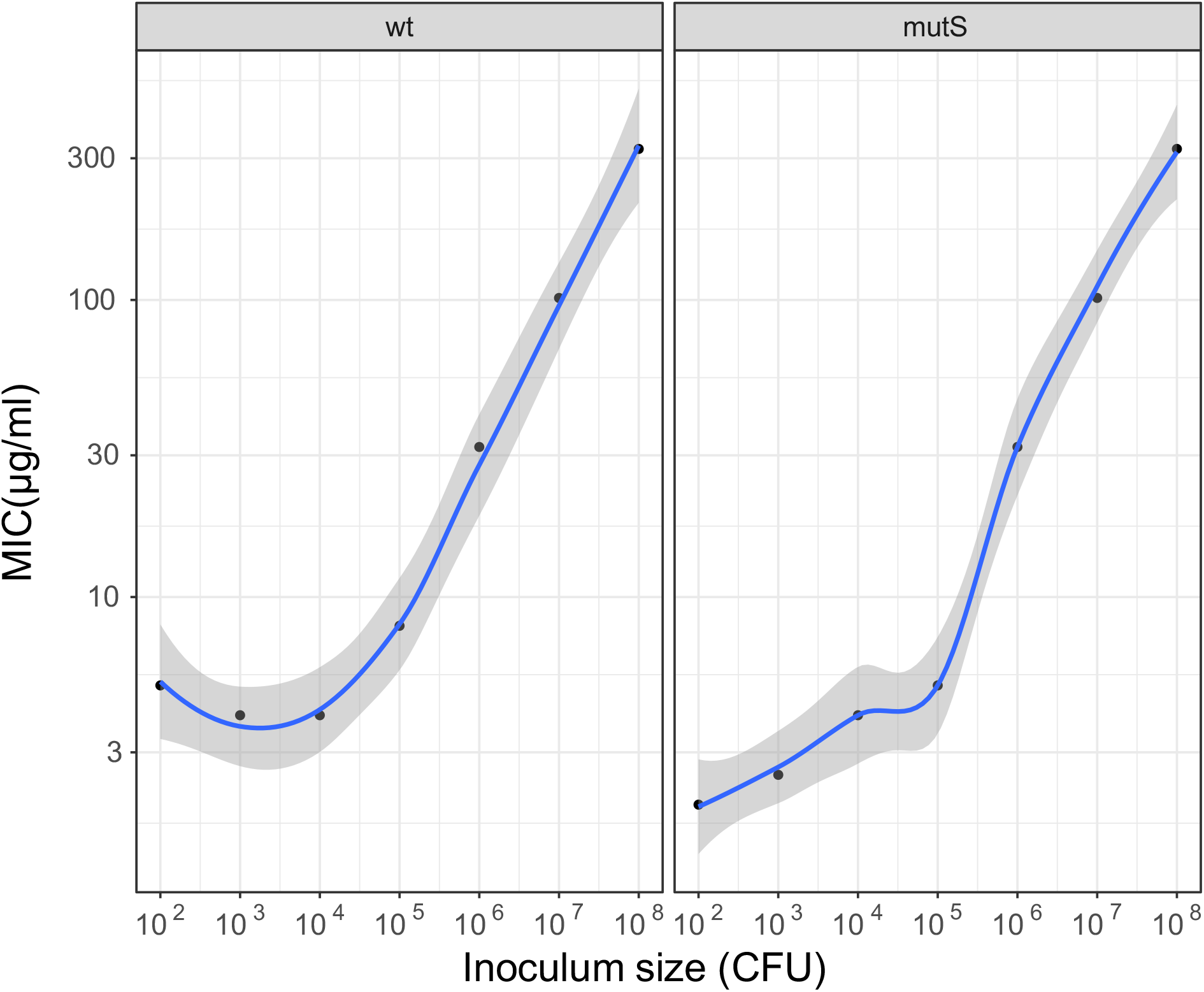
The effect of inoculum sizes for the MIC values of pexiganan. Each dot represents an average of 3 technical replicates. The grey areas display a 95% confidence interval.

## DISCUSSION

Here, we found that bacterial population sizes that are comparable to the *G. mellonella* gut during metamorphosis can influence the evolution of resistance of *E. coli* MG1655 to an AMP. This, however, was only realized if the mutation supply was high enough because only the mutator strains evolved resistance in our experiment.

Resistance against pexiganan only evolved in the *mutS* strain with an elevated mutation rate for both bottleneck sizes (**Figure 2b**). As expected, the highest resistance level occurred in the large *mutS* bottleneck due to the elevated mutation rate and population size combined. Although small *mutS* populations adapted to pexiganan, they did not increase their MIC as much as large *mutS* populations. This could have been caused by the reduced supply of beneficial mutations, which weaken selection for high-fitness individuals and enhance genetic drift favouring the accumulation of deleterious mutants (26). In the wt strain, we did not detect resistance evolution compared to the ancestor independent of the bottleneck size (**Figure 2b**) likely because of the insufficient mutation supply.

Theoretical work suggested that an increase in bottleneck size and mutation rate accelerates the adaptation. However, their combined effect on adaptation can be subtle. There is a growing number of studies on the adaptive potential of different bottleneck sizes (10,12,13,15–17). The majority of publications found that larger bottlenecks increase the adaptive potential (13,15– 17). A few cases based on the fitness landscape metaphor showed that smaller populations can out-compete large populations in adaptation to stressors (10,12). Additionally, *Wein et al. 2019*, showed that the effect of bottleneck size on adaptation can be context-dependent. Given that large bottlenecks were more advantageous than small bottlenecks for adaptation to higher temperatures (37 °C), however, there was no difference between bottleneck sizes for adaptation to 20 °C (29). Therefore, our study is consistent with the literature that reports that large bottlenecks provide an advantage to adapt better depending on the sufficient mutant supply.

Survival of bacterial populations and MIC assays provided different views on resistance evolution against pexiganan under different bottleneck sizes. The two large populations survived at AMP concentrations in the selection experiments, which are much higher than the measured final MICs (**Figure 2**). This disparity is most likely explained by the “inoculum effect”. The inoculum effect describes an increase in the MIC of a drug correlated with an increase in bacterial population size (30) (**Figure 3**): large populations have higher MICs than small populations. Therefore, in our experiment, small populations went extinct at lower concentrations than large populations. Thus, it is important to consider the inoculum effect to evaluate adaptation levels of different population sizes. Whether or not the inoculum effect plays out *in vivo*, for example, in the metamorphic insect gut or during an infection, is not known. In short, the MIC assay can be more informative than survival because, in the MIC assay all populations are inoculated with the same number of cells, which removes the inoculum effect.

Another reason that larger populations adapt better to antimicrobial selection could be that they, because of the inoculum effect, are exposed to sub-MIC concentrations for a longer time compared to small populations. This can increase the chance for beneficial mutations to emerge (31,32). Furthermore, in our experiments, large wt populations could not acquire sufficient beneficial mutations to improve their adaptation, whereas *mutS* (L) managed to adapt. Therefore, the interaction between bottleneck size and mutation supply seems important. It would be interesting to disentangle the effect of inoculum size from mutation rate on adaptation in future studies.

The different sizes of bottlenecks we studied here were informed by the bacterial population dynamics during metamorphosis in the moth *G. mellonella*. These bottlenecks are caused by the synthesis and release of antimicrobial peptides and lysozymes. AMPs are secreted markedly upon gut replacement at about stage V (fig. 1) (8). Therefore, we expected to observe the lowest microbiota count at stage V.

Can natural bottlenecks help insect hosts limit the emergence of AMP-resistant gut microbiota? Although our study showed that smaller bottleneck sizes evolved to lower MIC levels, our experimental design has some limitations. For instance, *E. coli* populations had prolonged exposure to pexiganan (e.g. 40 days for *mutS* large bottleneck size). By contrast, in natural conditions, *G. mellonella* gut replacement from stage 3 to 5 takes a few days, which provides a much shorter time window for potential resistance evolution (33). Yet, an *in vitro* study of *Staphylococcus aureus* demonstrated that resistance readily evolves against insect AMPs such as Tenecin 1 and Tenecin 2 in seven days (34). Although we only used *E. coli* as a proxy of the gut microbiota, in reality, the gut microbiota is more diverse. This can cause subtle interactions with the adaptation to the AMPs (9,22). Also, *G. mellonella* synthesizes a variety of AMPs that possibly synergize and reduce the chance of resistance (35).

All in all, the maintenance of gut microbiota can be important not only for insects but also other hosts. If hosts keep the gut microbiota abundance small, they could decrease the adaptive potential of, for example, pathogenic bacteria against AMPs. We speculate that such small population sizes as we observed here in insect guts and that we studied *in vitro*, are also a likely scenario for the start of infections in humans. Some work suggests that AMP resistance is a prerequisite for infections in humans (36). Such evolutionary dynamics during infection remain to be explored.

## Acknowledgments

We would like to thank Umut Değurmencu, Doğa Koçyiğit, Murat Tugrul, Ronan Murphy and Bernardo Antunes.

## SUPPLEMENTARY DATA

### Absolute MIC of treatments and experimental controls

### MIC fold change increase of *mutS* populations throughout the experiment

**Figure S1.**
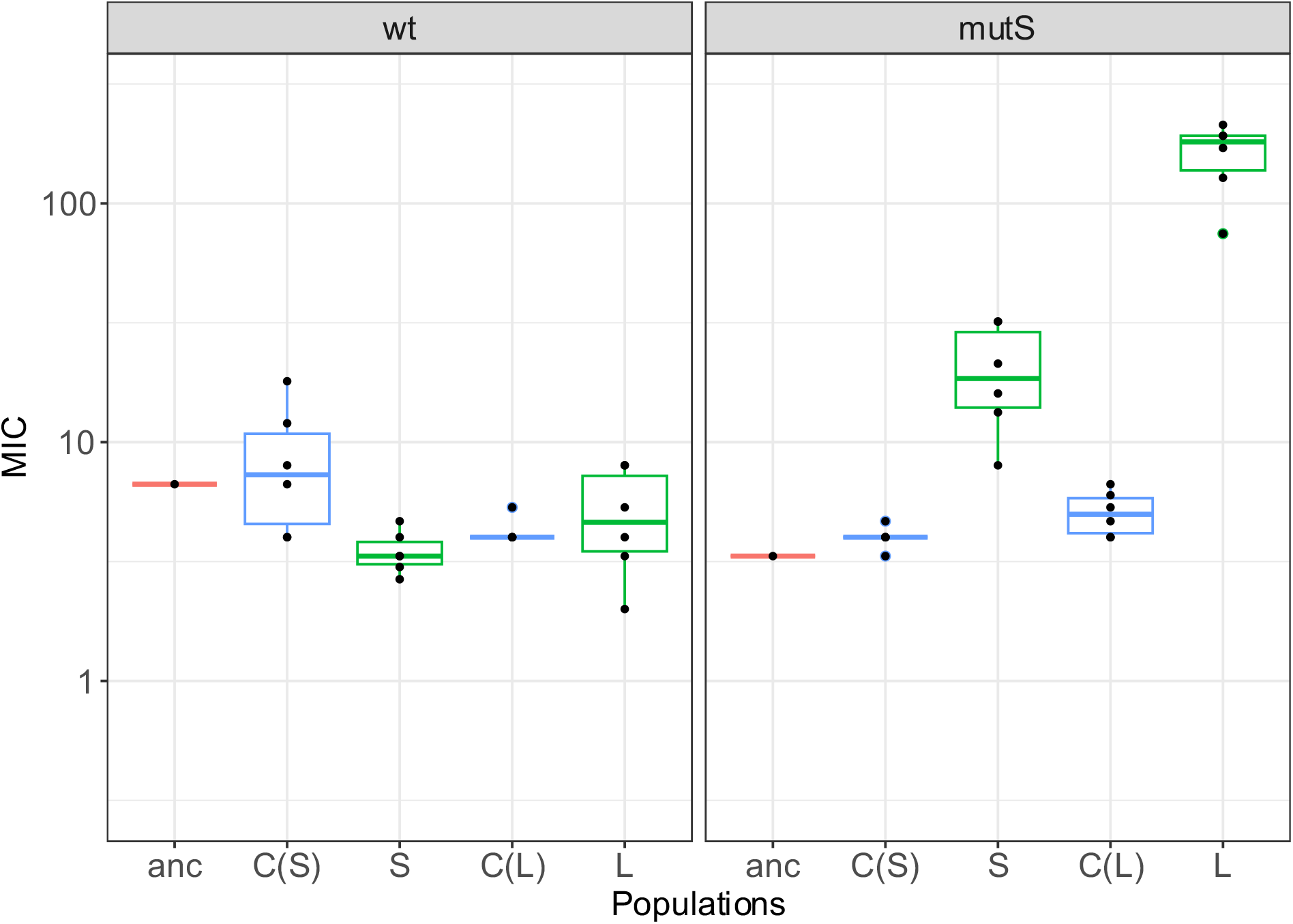
Absolute MIC values of ancestors (anc), experimental controls for small and large population sizes (C(S), C(L)), and treatments (S, L) for wildtype(wt) and mutator strains (mutS).

**Figure S2.**
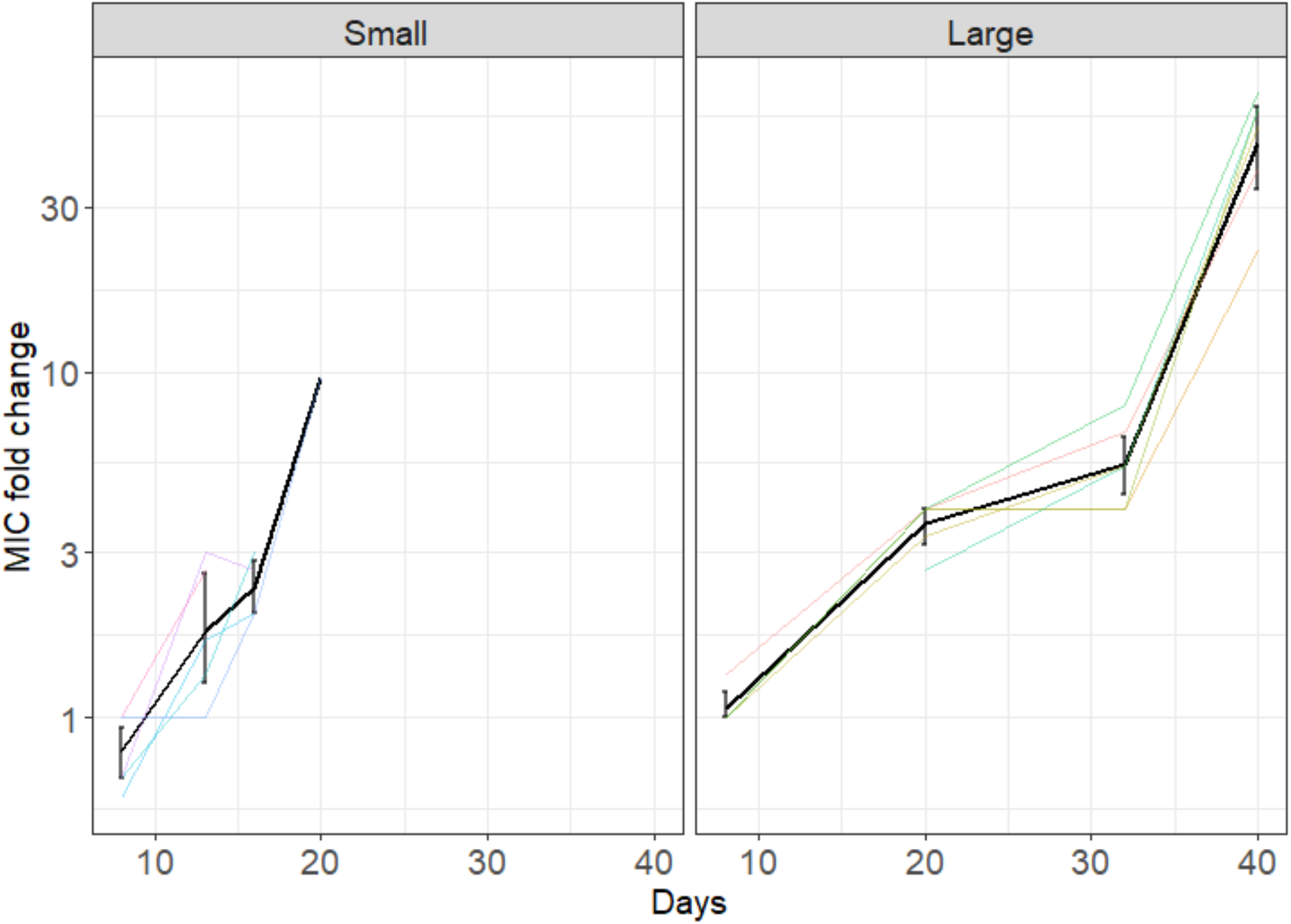
The MIC fold change of small and large bottlenecks of mutS during the selection experiment. The MIC of treatment populations was compared to their ancestor. Colored solid lines show each replicate and black solid lines with error bars (standard error) display the average MIC fold change across replicates.

**Table S1:** The comparison of gut microbiota abundance between different developmental stages(Tukey HSD).

|  | **difference** | **lower** | **upper** | **P adjusted** |
| --- | --- | --- | --- | --- |
| **1-0** | -0.227 | -0.898 | 0.444 | 0.88 |
| **3-0** | -0.891 | -1.596 | -0.187 | 0.006 |
| **5-0** | -0.999 | -1.722 | -0.276 | 0.002 |
| **A-0** | 0.199 | -1.197 | 1.596 | 0.995 |
| **3-1** | -0.665 | -1.327 | -0.003 | 0.049 |
| **5-1** | -0.772 | -1.453 | -0.091 | 0.018 |
| **A-1** | 0.426 | -0.949 | 1.802 | 0.909 |
| **5-3** | -0.108 | -0.822 | 0.607 | 0.993 |
| **A-3** | 1.091 | -0.302 | 2.483 | 0.196 |
| **A-5** | 1.198 | -0.203 | 2.6 | 0.13 |

